# Functional Brain Networks and Alcohol Consumption: From the Naïve State to Chronic Heavy Drinking

**DOI:** 10.1101/2020.09.14.296871

**Authors:** Jared A. Rowland, Jennifer R. Stapleton-Kotloski, Greg E. Alberto, April T. Davenport, Phillip M. Epperly, Dwayne W. Godwin, James B. Daunais

## Abstract

A fundamental question for alcohol use disorder is how naïve brain networks are reorganized in response to the consumption of alcohol. The current study aimed to determine the progression of alcohol’s effect on functional brain networks during the transition from naïve, to early, to chronic consumption. Resting-state brain networks of six female monkeys were acquired using magnetoencephalography prior to alcohol exposure, after early exposure, and after free-access to alcohol using a well-established model of chronic heavy alcohol use. Functional brain network metrics were derived at each time point. Assortativity, average connection frequency, and number of gamma connections changed significantly over time. All metrics remained relatively stable from naïve to early drinking, and displayed significant changes following increased quantity of alcohol consumption. The assortativity coefficient was significantly less negative (*p*=.043), connection frequency increased (*p*=.03), and gamma connections increased (*p*=.034). Further, brain regions considered hubs (*p*=.037) and members of the Rich Club (*p*=.012) became less common across animals following the introduction of alcohol. The minimum degree of the Rich Club prior to alcohol exposure was significantly predictive of future free-access drinking (r=-.88, *p*<.001). Results suggest naïve brain network characteristics may be used to predict future alcohol consumption, and that alcohol consumption alters the topology of functional brain networks, shifting hubs and Rich Club membership away from previous regions in a non-systematic manner. Further work to refine these relationships may lead to the identification of a high-risk AUD phenotype.

## Introduction

Alcohol use disorder (AUD) constitutes a global problem and is ranked among the top substance abuse problems in the United States, with over 70% of adults that struggle with substance use disorder estimated to abuse alcohol ^1^(SAMSHA 2018). AUD impacts global brain functional networks including the default mode, executive, attentional, salience and reward networks ^2–7^ but the neurocircuitry underlying vulnerability and resilience to alcohol use disorder (AUD) is not clearly understood, making it difficult to establish viable, targeted treatment options.

This lack of clarity is due, in part, to the difficulty in capturing an alcohol-naïve baseline in human subjects. Clinical studies are often conducted with long-term drinkers at different drinking phases after changes in brain networks have likely already manifested. It is clear that AUD is characterized in part by dysfunctional information processing ^8^ that occurs in part through altered brain activity during both resting state (RS) and task performance in alcoholics ^9^ as compared to other neurological conditions ^10^. Functional brain networks including the default mode, salience, and executive networks are known to be sensitive to chronic alcohol use ^11,12,7^ however, the temporal nature and anatomic directionality of changes that occur remains unclear.

A limitation of studies involving humans is the significant individual variation in history regarding substance use, comorbid exposure to substances, and living environments. NHP models ^13^ have demonstrated that daily drinking for 15 months causes functional and genomic changes across the brain when contrasted against the alcohol naïve brain ^14–21^, as well as reorganization of brain networks measured by fMRI ^22^ and significantly altered signal power of multiple bandwidths across the brain using magnetoencephalography (MEG) ^23^.

The objective of the current study was a longitudinal examination of the trajectory of these changes to identify at what point they begin to manifest and if baseline, alcohol-naïve indicators of future drinking can be identified.

## Materials and Methods

### Animals

Adult female rhesus monkeys (n=6, 5-7 years old at study start) were subjects in an ongoing ethanol (EtOH) self-administration study. This age group reflects late adolescence to early adulthood in humans. The monkeys were trained on an operant panel to self-administer all fluids and food using a well-established drinking model ^13,24^. This process begins with EtOH-naïve monkeys that are induced to drink escalating doses of EtOH (0.5, 1.0 and 1.5 g/kg) for 30 days at each dose (induction phase). All monkeys were maintained at 1.5 g/kg for 20 drinking days while operant panels were serviced and re-programed. Animals were then provided free access to EtOH and water for 22 hours per day, 5 days per week for 180 days. Sessions began at 11:00 am each day. MEG recordings were acquired under EtOH naïve conditions (Baseline), after completion of the induction phase (Post Induction) and after 180 open access drinking days (Free Access). Control animals were not utilized in the current study due to the previously well-established effects of EtOH in this model contrasting EtOH-exposed and control animals ^14–21,23^.

### Preparation for MEG scans

Animals were fasted overnight from food but not EtOH prior to scans. On the day prior to the Post-Induction scan all animals had consumed the provided 1.5 g/kg EtOH dose by 6 pm. The time to finish the 1.5 g/kg dose ranged from 6-300 minutes. Average time between last drink and sedation for imaging was 1183.0 minutes (SD=124.7, min=1025, max=1340) at the Post-induction scan and 344.2 minutes (SD=377.9, min=0.0, max=977) at the Free Access Scan. These time frames raise the possibility of acute withdrawal ^25^; however, symptoms were not observed during similar time frames on non-imaging days ^26^ and the anesthetic agent (propofol, a GABAa receptor positive allosteric modulator ^27^) helped ensure acute withdrawal symptoms were not present during data acquisition. Previous work has shown that acute withdrawal in this model peaks between 24 - 72 hours ^20^, which is beyond the duration of abstinence present here. Animals were sedated with ketamine (12 mg/kg, i.m.) for transport to the MEG suite. Anesthesia was induced with a bolus injection of 2.0-4.0 mg/kg propofol to allow intubation and was maintained via intravenous continuous infusion of 200 μg/kg/min propofol via syringe pump (Sage, Orion Research Corporation, Cambridge, Mass). Animals were placed in a supine position and artificially ventilated. These preparations are consistent with our previous reports ^22,23^.

### Magnetoencephalography recordings

Data were acquired using a whole head CTF Systems Inc. MEG 2005 neuromagnetometer system equipped with 275 first-order axial gradiometer coils. Head localization was achieved using a conventional three-point fiducial system (nasion and preauricular points). Each monkey was tattooed at each fiducial location to ensure consistent placement over time. Resting-state recording was conducted with animals lying supine for 5 minutes. Data were sampled at 1200 or 2400 Hz over a DC-300 or DC-600 Hz bandwidth, respectively. MEG data were preprocessed using synthetic 3^rd^ order gradient balancing, whole trial DC offset, and band pass filtered from DC-80 Hz with powerline filtering. Data were visually inspected for obvious muscle artifact, and such epochs, if present, were discarded from further analyses. Following initial MEG recording, a T1 weighted MRI image was obtained for each animal for co-registration and localization of MEG signals.

### Network analysis

Network analysis was conducted identically to previous work ^28,29^. Network analysis proceeded by first identifying nodes of the network and quantifying communication among those nodes. The resulting matrices are conducive to the application of graph theory for calculating metrics describing the topology of the network.

### Network creation

For each animal 41 non-adjacent bilateral regions of interest (ROIs, 2 mm^3^) were identified in native brain space. ROIs were chosen to represent the default mode and reward networks. These networks have been previously demonstrated to be affected by chronic heavy alcohol consumption in humans ^3,30,31^. Brain regions included the anterior cingulate, medial and lateral orbital frontal cortex, principle sulcus, nucleus accumbens, caudate head and body, head of the putamen, parietal area, precuneus, lateral and medial amygdala, anterior, medial, and posterior hippocampus, vermis, anterior and posterior lobes of the cerebellum, thalamus, and anterior insula. Source series representing the unique weighted sum of the output across all MEG sensors for a specific ROI in the brain were calculated using a well-validated beamformer (synthetic aperture magnetometry, SAM) ^32,33^. The weighted phase lag index (wPLI); ^34^ was calculated between all pairs of nodes using the source series to measure functional connectivity, filtered between 1 and 80 Hz. The wPLI is a phase-based metric insensitive to fluctuations in source amplitude. Connectivity was operationalized at the frequency with the highest wPLI value, allowing the frequency at which connections occurred to vary from connection to connection. This represents a better model of brain activity than restricting connectivity to a specific frequency band. Data were first thresholded using 5,000 unique pairs of phase-randomized surrogate time series calculated for each animal individually (Prichard and Theiler, 1994) to remove connections not different from noise. The resulting networks were then thresholded by satisfying the equation S=log(N)/log(K) where N represents the number of nodes in the network and K the average degree using S=2.5 ^35^.

### Network metrics

Network metrics calculated are listed in Table 1. Metrics were selected with a focus on characterizing the topology of the overall network. *Clustering Coefficient* was selected as an indicator of clustering and subgroup formation within the network. This metric was calculated as defined in Stam, Reijneveld ^36^. *Modularity* was selected as an indicator of well-defined subnetworks within the larger network. This metric was calculated using the Louvain method of community detection as defined in Blondel, Guillaume, Lambiotte, Lefebvre ^37^. The analysis was run 500 times, using the average number of modules (*Number Modules*) as outcome variables. *Assortativity coefficient* represents the correlation coefficient of the degree of nodes on each end of a connection. The degree of a node is the number of direct connections that node has to other nodes in the network. A positive coefficient suggests nodes are preferentially connecting to other nodes of similar degree, while a negative coefficient suggests nodes preferentially connect to those of different degree ^38^. Rich Club was selected as an indicator of the presence of a “network backbone”. The Rich Club is a subset of highly connected and highly interconnected nodes forming the basis of the broader network. Rich Club metrics were calculated as defined in Colizza, Flammini, Serrano, Vespignani ^39^ using 500 independently generated random networks. The number of nodes (Rich Club Nodes) within the Rich Club, the minimum degree of those nodes (Rich Club Degree), and interconnectivity among those nodes (Rich Club Coefficient) were used as outcome variables. The Rich Club Coefficient was weighted by the average of the same metric across the 500 random networks, representing the level of increased interconnectivity over a random network. *Hubs* of the network were identified as the 10% of nodes (*n*=4) with the highest degree.

**Table 1.** Descriptive statistics of network metrics prior to and following exposure to ethanol.

| Network Metric | Baseline | Post-Induction | Free Access |
| --- | --- | --- | --- |
| Clustering Coefficient, mean (SD) | 0.35(0.1) | 0.28(0.1) | 0.30(0.1) |
| Assortativity coefficient, mean (SD)<sup>a,b</sup> | -0.40(0.1) | -0.44(0.1) | -0.19(0.2) |
| Rich Club Coefficient, mean (SD) | 1.84(0.1) | 1.72(0.1) | 1.91(0.3) |
| Rich Club Nodes, mean (SD) | 13.17(1.7) | 14.5(2.9) | 10.83(4.3) |
| Rich Club Minimum Degree, mean (SD) | 7.50(1.1) | 6.67(0.8) | 9.50(2.7) |
| Number of Modules, mean (SD) | 10.50(3.4) | 7.83(2.4) | 8.83(4.4) |
| Mean Connection Frequency, mean (SD)<sup>a,c</sup> | 6.57(5.7) | 5.61(4.9) | 16.01(10.6) |
| Mode Connection Frequency, mean (SD) | 4.39(5.9) | 4.79(6.0) | 11.13(14.3) |
| Delta Connections, mean (SD) | 172.67(96.0) | 202.0(79.5) | 99.0(99.7) |
| Gamma Connections, mean (SD)<sup>a</sup> | 1.67(4.1) | 4.0(8.9) | 54.67(64.5) |

Note. $n = 6$ , <sup>a</sup>Omnibus test of change over time $p < .05$ , <sup>b</sup>paired samples t-test between Post Induction and 6 months $p < .05$ ; <sup>c</sup>paired samples t-test between EtOH Naïve and 6 months $p < .05$ ; EtOH = ethanol, SD = Standard Deviation, follow-up paired samples t-tests were only conducted when the Omnibus test was significant.

### Materials

Beamforming and source series construction were completed using software provided by CTF MEG International Services LP (Coquitlam, BC, Canada). Further analyses of source series data and network creation were conducted using Matlab 2016a. Network metrics were calculated using the Brain Connectivity Toolbox ^40^. SAS Enterprise Guide 7.1 (SAS Institute Inc., Cary, NC) was used for statistical analysis.

### Analyses

Differences across time in network metrics (3 time-points) were examined using repeated-measures ANOVA. When the omnibus effect of Time was significant, additional paired sample t-tests were used to identify the differences among specific time points. Consistency in hubs and Rich Club membership were examined using the number of animals for which a region was considered a hub or Rich Club member as the dependent variable and brain regions listed in Tables 3 and 4 as the independent variable in a repeated measures ANOVA. Brain regions were further characterized as highly common (> 3 animals) or not. Spearman rank correlations were conducted to examine the relationship between network metrics and drinking outcomes (daily average g/kg). Two-tailed tests and alpha of 0.05 were used for significance.

**Table 2.** Baseline network metrics correlated with Free Access drinking levels.

|  | Conn<br>Freq | Rich Club<br>Coefficient | Rich Club<br>Nodes | Rich Club<br>Degree | Clustering<br>Coefficient | Assortativity |
| --- | --- | --- | --- | --- | --- | --- |
| Free Access Avg<br>Daily g/kg | 0.60 | -0.26 | 0.71 | -0.88<sup>a</sup> | -0.31 | 0.03 |

Note. $n = 6$ , Free Access Avg Daily g/kg = daily consumption in g/kg during the 180 days of unrestricted access, Conn Freq = Connection Frequency, Avg = Average, g = gram, kg = kilogram, <sup>a</sup> $p = .02$ .

**Table 3.** The number of subjects for which each brain region was considered a hub of the network. Nodes are considered hubs if they in the upper 10% of the network for degree. Only node considered a hub for at least 2 subjects at any time point are presented.

| Brain Region | Baseline | Post-Induction | Free Access |
| --- | --- | --- | --- |
| Parietal | 3 | 2 | 0 |
| Thalamus | 4 | 2 | 2 |
| Precuneus | 2 | 1 | 1 |
| Cerebellum | 4 | 4 | 2 |
| Amygdala | 2 | 0 | 2 |
| Hippocampus | 2 | 2 | 4 |

Note. $n = 6$ .

**Table 4.** The number of subjects for which each brain region was considered a member of the Rich Club.

| Brain Region | Baseline | Post-Induction | Free Access |
| --- | --- | --- | --- |
| Putamen | 5 | 5 | 4 |
| Hippocampus | 5 | 6 | 5 |
| Thalamus | 5 | 5 | 3 |
| Insula | 5 | 5 | 2 |
| Cerebellum | 5 | 6 | 3 |
| Parietal | 4 | 4 | 3 |
| Amygdala | 4 | 4 | 5 |
| OrbitoFrontal | 4 | 4 | 3 |
| Caudate | 3 | 6 | 3 |
| Precuneus | 3 | 1 | 2 |

Note. $n = 6$ .

## Results

Daily average g/kg EtOH consumption (see Figure 1) increased significantly once given free access (mean [SD]; Post-Induction=1.0 [0.0], Free-Access=4.7 [1.0], *p*<.01, Cohen’s d=5.23). Network metrics at Baseline, Post-Induction (after 120 days escalating controlled doses), and Free-Access (after 180 days unrestricted access) are shown in Table 1.

**Figure 1.**
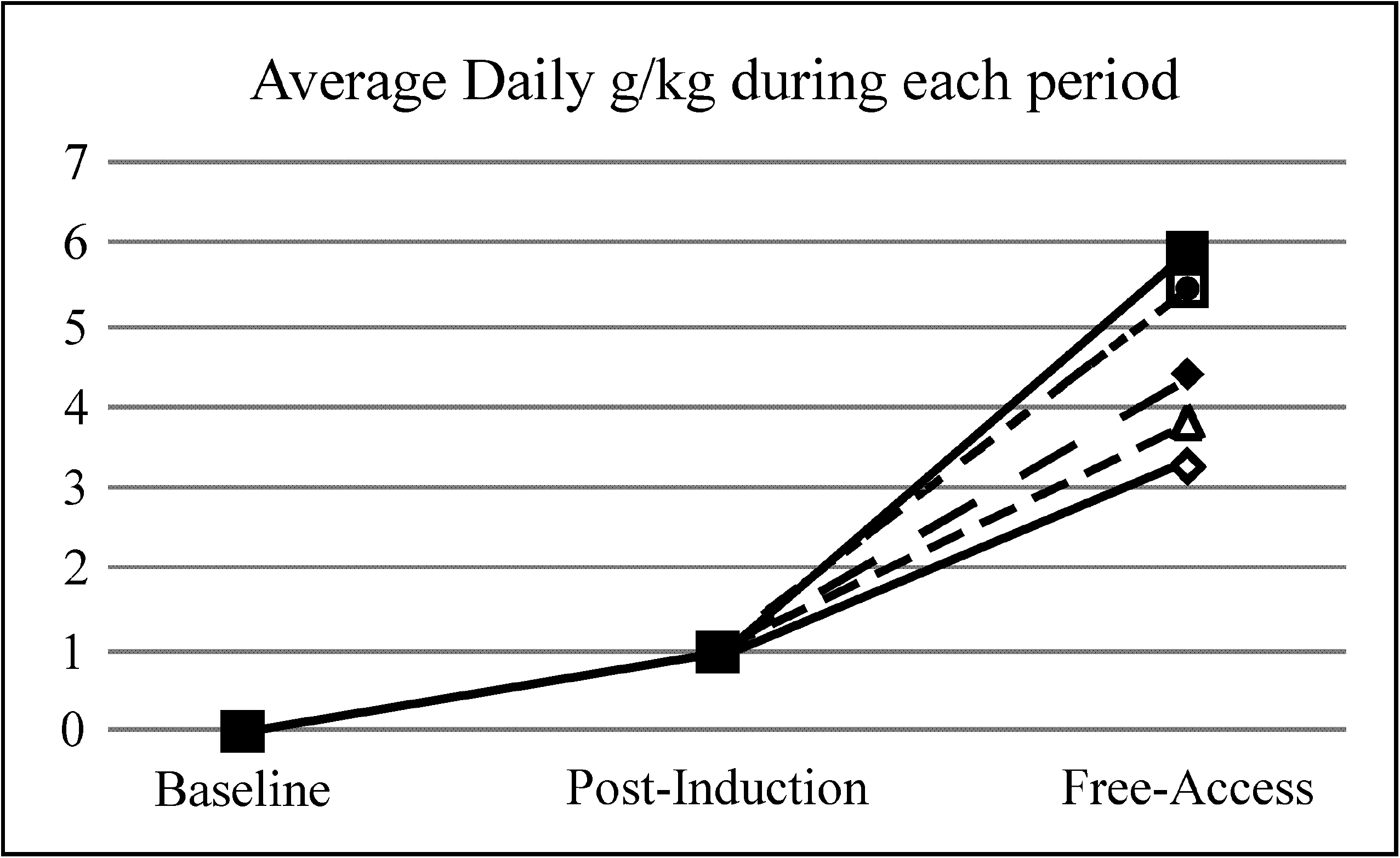
Average daily g/kg EtOH consumption during the Baseline (naïve), Induction (0.5, 1.0, and 1.5 g/kg at 30 day intervals), and Free Access (180 days unrestricted) periods.

### Networks predicting EtOH consumption

Table 2 illustrates correlations between Baseline (EtOH naïve) network metrics and Free Access consumption. The minimum degree of the Rich Club at baseline was strongly related to Free Access consumption levels (Figure 2). No correlation between Post-Induction network metrics and Free Access EtOH consumption reached significance, thought the qualitative pattern was similar to Baseline results.

**Figure 2.**
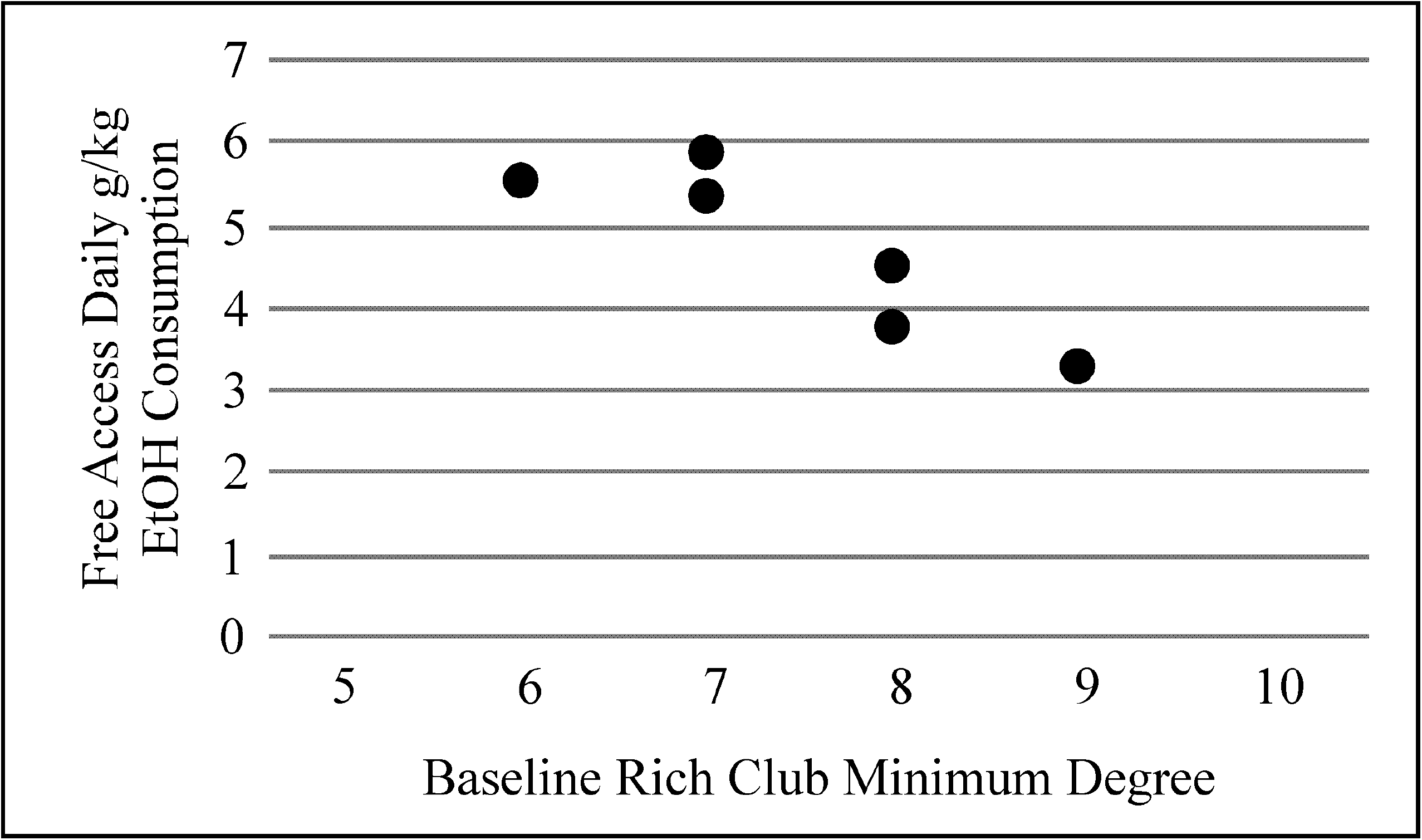
The minimum degree of the Rich Club of the alcohol naïve (Baseline) functional brain network was strongly related to future drinking when animals were provided free-access to alcohol (r=−.88, *p*=.02).

### Effects of EtOH on networks

Assortatitvity coefficient, F(2,8)=4.77, *p*=.043), average connection frequency, F(2,8)=5.64, *p*=.030, and the number of connections in the gamma bandwidth, F(2,8)=5.33, *p*=.034 changed significantly over time (see Table 1). The assortativity coefficient was negative and stable from baseline to post-induction, suggesting disassortivity, but increased significantly following chronic exposure to EtOH (contrasting Post-Induction against Free Access, t(5)=-3.63, *p*=.015). The average connection frequency remained stable from Baseline to Post-Induction, was significantly higher at Free Access than at Baseline, t(5)=-3.20, *p*=.024, but not Post Induction, t(5)=-2.48, *p*=.056. Finally, while the omnibus test of change in the number of connections in the gamma bandwidth over time was significant, no post-hoc contrast reached statistical significance. Qualitatively, the number of connections increased upon Free Access, mirroring the previous two findings.

### Specific Brain Regions

Table 3 demonstrates areas considered hubs across animals at each time point. For brain structures with multiple aspects, the region was considered a hub if any of the aspects were considered a hub (e.g. if either the posterior, medial, or anterior hippocampus was a hub, then Table 3 indicates the hippocampus as a hub). There was an interaction between brain regions and time, F(2,10)=4.67, *p*=.037, such that those brain regions commonly serving as hubs across animals (i.e. thalamus and cerebellum) at baseline became less so following the introduction of alcohol. Table 4 includes brain regions that were members of the Rich Club for at least three animals at any time point, again collapsing within regions. A significant effect of time was observed, F(2, 16)=5.91, *p*=.012 demonstrating a decrease in the commonality of regions in the Rich Club across animals. There was no interaction with common regions at baseline.

## Discussion

The current study demonstrates that aspects of the Rich Club of alcohol-naïve brain networks are strongly related to future drinking behaviors. In addition, results demonstrate increasing alterations of network topology as the duration of exposure to alcohol increases and the quantity of alcohol consumed increases. As exposure and quantity increased, the hubs of the network and membership of the Rich Club were observed to shift, becoming less consistent across animals.

Aspects of *alcohol-naïve* resting-state functional brain networks were demonstrated to predict *future* drinking levels. The minimum degree of the Rich Club observed at Baseline (alcohol-naïve) was strongly and inversely correlated with the level of alcohol consumption during the future free access period. Interconnectivity among the Rich Club nodes (Rich Club Coefficient) was not related to future drinking, demonstrating that as the interconnectivity increased between the Rich Club members and the rest of the network at baseline, future alcohol consumption decreased. This suggests that Rich Club characteristics of functional brain networks may be predictive of future drinking levels, *even when measured prior to alcohol exposure*.

These results are consistent with previous work using the same NHP model indicating that premorbid behaviors and those occurring early in the drinking history may be predictive of future consumption levels. These factors include low cognitive flexibility ^41^, early drinking phenotypes (i.e., gulping vs sipping) ^13,42^, age, latency to begin drinking, and the number of “bouts” of drinking ^42,43^. The current results are the first to provide a brain-based factor indicative of future drinking in this model, suggesting that premorbid alcohol-naïve differences in Rich Club characteristics of functional brain networks also predict future drinking in this model.

These results extend recent findings in human participants demonstrating that white matter brain networks of individuals with AUD displayed lower Rich Club characteristics compared to their non-abusing siblings, who displayed lower levels compared to control participants ^44^. While Zorlu, Capraz, Oztekin, Bagci, Di Biase, Zalesky, Gelal, Bora, Durmaz, Besiroglu, Saricicek ^44^ suggest potential premorbid differences in white matter network structure may be a marker of risk or susceptibility to AUD, the results of the current study provide direct empirical support for this hypothesis, showing that Rich Club characteristics of premorbid functional brain networks are directly related to future drinking levels.

Alcohol-induced changes in functional brain networks were observed following significant increases in the quantity of alcohol consumed. Networks remained disassortive, but significantly less so. In addition, the mean connection frequency increased significantly, mirrored by a non-significant increase in the mode of the connection frequency. There was a significant increase in the number of connections in the gamma bandwidth mirrored by non-significant decreases in the number of connections in the delta bandwidth. Significant changes were also seen in the backbone of networks. Brain regions considered hubs and members of the Rich Club were more common across animals at baseline than following the introduction of alcohol, suggesting the networks were being altered in an inconsistent manner across animals. It should be noted that the quantity of alcohol consumed during the induction period is relatively low (0.5 to 1.5 g/kg) and increased between 100% and 300% during unrestricted access. These results support the potential for a dose-dependent relationship between patterns of alcohol consumption and the structure of functional brain networks ^45–47^.

Limitations of the current study include the small sample size, which limits the complexity and sensitivity of analyses that can be conducted. Neuroimaging was conducted under anesthesia, which has known effects on brain function ^48–50^. Possible interactions between the anesthetic and alcohol could have occurred, if not directly, then through the indirect development of tolerance. However, anesthesia was maintained at consistent levels and physiological indicators of arousal were monitored continuously, suggesting that levels of sedation were consistent across scans. Neuroimaging under conscious conditions will be required to completely understand the effects of alcohol on brain function using this model. The interval between ethanol access and MEG scans raises the possibility that some animals may have been experiencing symptoms of withdrawal ^25^. However, signs of withdrawal were not observed during the same time periods on non-imaging days ^20,26^. Also, propofol was used as the anesthesic agent, helping to ensure animals were not experiencing withdrawal symptoms during scans ^27^. Finally, animals who ceased alcohol consumption prior to scans did so voluntarily and in a time frame consistent with non-imaging days and were not forcibly fasted.

The Rich Club is a community of nodes within the network that have high degree and interconnectedness ^39^. These nodes represent hubs within the network and communication among these nodes can often serve as “shortcuts” within the network, increasing efficiency of communication across otherwise distantly connected nodes. Alterations to this important subnetwork are likely to have broad and sweeping effects on brain communication and information processing ^51^. However, differences in Rich Club characteristics have been observed in many neurodevelopmental disorders, including schizophrenia ^52^, bipolar disorder ^53^, and autism ^54^. As such, the broad differences in Rich Club characteristics observed in this study are unlikely to serve as a direct “neurophenotype” of alcohol use disorder without further refinement and empirical study. However, these results identify that differences in network topology are important to understanding individuals who might be at risk for future heavy drinking or AUD.

Assortativity describes the tendency for connectivity to occur between nodes similar in a particular characteristic, usually degree. The assortativity coefficient was strongly negative at baseline, suggesting disassortativity, or the tendency for nodes to connect to other nodes of *different* degree. This is common in biological networks and suggests a hierarchical nature to connectivity supportive of clustering and modularity around central hubs^38^. As demonstrated by the current results, increased quantity of alcohol consumption disrupted the disassortativity of networks. This is further supported by reduced commonality in network hubs (Table 3) and Rich Club membership (Table 4) across time.

## Conclusions

The current study identified a relationship between functional brain networks in the alcohol-naïve state and future alcohol consumption, consistent with other work using this model demonstrating early behavioral markers of future drinking. Additionally, significant alterations to the topology of the network were observed following the significant increase in quantity of alcohol consumption during the Free Access period. Future work will be invaluable in clarifying the changes, and potentially the timing of those changes, that infer risk specific to AUD in humans.

## Author Contributions

Conception and study design (JBD), data collection and acquisition (JBD, JAR, JRS, ATD, GEA, PME), statistical analysis (JAR, JRS), interpretation of results (JBD, JAR, JRS, DWG), drafting the manuscript or revising it critically for important intellectual content (All Authors), and approval of final version to be published and agreement to be accountable for the integrity and accuracy of all aspects of the work (All authors).

## Acknowledgements

This study was funded through P01 AA021099-S1 (JBD), P50 AA026117 (JBD, DWG, JAR) AA016852 (DWG), and 5F30AA023708-02 (GEA), Ignition Funds from the Wake Forest School of Medicine Translational Sciences Institute (JBD) and a TSI pilot award (DWG) and support from the Department of Neurology, the Center for Biomolecular Imaging, the Wake Forest University Primate Center, the W.G. (Bill) Hefner Veterans Affairs Medical Center and VA Mid-Atlantic Mental Illness, Research, Education, and Clinical Center.

## Compliance with Ethical Standards

This work was approved by the Wake Forest School of Medicine Institutional Animal Care and Use Committee and conducted in compliance with all regulations, including measures taken to reduce pain and suffering.

